# Accurate detection of convergent amino-acid evolution with PCOC

**DOI:** 10.1101/247296

**Authors:** Carine Rey, Laurent Guéguen, Marie Sémon, Bastien Boussau

## Abstract

In the history of life, some phenotypes have been acquired several times independently, through convergent evolution. Recently, lots of genome-scale studies have been devoted to identify nucleotides or amino acids that changed in a convergent manner when the convergent phenotypes evolved. These efforts have had mixed results, probably because of differences in the detection methods, and because of conceptual differences about the definition of a convergent substitution. Some methods contend that substitutions are convergent only if they occur on all branches where the phenotype changed towards the exact same state at a given nucleotide or amino acid position. Others are much looser in their requirements and define a convergent substitution as one that leads the site at which they occur to prefer a phylogeny in which species with the convergent phenotype group together. Here we suggest to look for convergent shifts in amino acid preferences instead of convergent substitutions to the exact same amino acid. We define as convergent shifts substitutions that occur on all branches where the phenotype changed and such that they correspond to a change in the type of amino acid preferred at this position. We implement the corresponding model into a method named PCOC. We show on simulations that PCOC better recovers convergent shifts than existing methods in terms of sensitivity and specificity. We test it on a plant protein alignment where convergent evolution has been studied in detail and find that our method recovers several previously identified convergent substitutions and proposes credible new candidates.

## Introduction

Convergent phenotypic evolution provides unique opportunities for studying how genomes encode phenotypes, and for quantifying the repeatability of evolution. These questions are typically addressed by sequencing genes or genomes belonging to a sample of species sharing a convergent phenotype, along with those of closely related species sharing a different ancestral phenotype. Then, nucleotide or amino acid positions that are inferred to have changed specifically on those branches where the phenotypes convergently changed may be assumed to be involved in the convergent evolution of those phenotypes. Such an approach has been used on spectacular cases of convergent evolution such as the C4 metabolism in grasses (Besnard *et al*., 2009), the ability to consume a toxic plant compound in insects (Zhen *et al*., 2012), echolocation in whales and bats (Parker *et al*., 2013), or the ability to live in an aquatic environment in mammals (Foote *et al*., 2015). These studies have found different levels of convergent evolution. In particular Parker *et al*. (2013) investigated convergent substitutions associated with the evolution of echolocation in mammals, which has evolved once in whales and once or twice in bats. They focused on amino acid sequences rather than on nucleotide sequences, assuming that it is where most selective effects would be observed. Using a topology-based method, they found a large number of convergent substitutions in close to 200 genes. However when these protein data were reanalyzed using another method, it was concluded that many of those convergent changes were likely false positives (Thomas and Hahn, 2015; Zou and Zhang, 2015b).

These strong disagreements come from differences in the bioinformatic methods that were used to detect convergent substitutions, and the underlying definition of what makes a substitution convergent. If we put aside studies of individual genes that involved manual analyses of alignments and detailed investigations of the rate of sequence evolution and patterns of selection along gene sequences (Besnard *et al*., 2009; Zhen *et al*., 2012), genomic studies have relied on two different methods. In (Zhang and Kumar, 1997), and later in (Foote *et al*., 2015; Thomas and Hahn, 2015; Zou and Zhang, 2015b), convergent sites are defined as those that converged to the exact same amino acid in all convergent species. Instead, in (Parker *et al*., 2013), a more operational definition is used: a convergent site is one that prefers to the species phylogeny a phylogeny in which species with the convergent phenotype group together. In doing so, they have no explicit requirement over the type of amino acid change that occurred in the species with the convergent phenotype because their method is remote from the actual mechanism of substitutions. With a more relaxed definition than in (Thomas and Hahn, 2015; Zou and Zhang, 2015b), it is not surprising that they recover more instances of convergent amino acid evolution.

### From convergent substitutions to convergent shifts

We believe that these two definitions have several shortcomings. First, the historical definition of (Zhang and Kumar, 1997) seems very strict. Selecting only sites that converged to the exact same amino acid in all species with a convergent phenotype is bound to capture only a subset of the substitutions associated with the convergent phenotypic change. This will capture only those sites where a unique amino acid is much more fit in the convergent phenotype than all other amino acids. In many other cases, there may be more than one amino acid that is fit at a particular position, given the convergent phenotype. For instance, it may be that several amino acids with similar biochemical properties have roughly the same fitness at that site. In such circumstances, we do not expect that identical amino acids will be found in all species with the convergent phenotype, but that several amino acids with similar biochemical properties will be found in all species with the convergent phenotype. Such convergent shifts in the amino acid preference at a given site are not considered under the definition of (Foote *et al*., 2015; Zhang and Kumar, 1997). Second, (Parker *et al*., 2013)’s definition may be too loose, as it is entirely disconnected from the substitution process.

We propose to consider shifts in amino acid preference instead of convergent substitutions. To us, a substitution is convergent if it occurred towards the same amino acid preference on every branch where the phenotype also changed towards the convergent phenotype. We model the amino acid preference at a position and on a branch by a vector of amino acid frequencies, which we call a profile. The amino acid profile used in species with the convergent phenotype needs to be different from the profile used in species with the ancestral phenotype. This definition conveys the idea that a convergent substitution is necessary to a convergent phenotype, that is, every time the phenotype changes to the convergent state, the position must change towards the convergent phenotype. It is thus equivalent to (Zhang and Kumar, 1997)’s definition in its positioning of changes on the branches where the phenotypic change occurred, but it seems less restrictive from a biochemical point of view. It extends previous works (Parto and Lartillot, 2017, 2018; Studer *et al*., 2014; Tamuri *et al*., 2009) that also modeled changes in amino acid profiles, but did not require that there should be a change on the branch where the phenotype changed from ancestral to convergent.

### Detecting convergent shifts

In this manuscript, we evaluate our proposed definition by comparing a method that uses our definition to two other methods proposed in the literature to detect convergent substitutions.

The power of a method is usually analyzed in terms of specificity and sensitivity. Specificity is critical for methods that detect convergent substitutions. Specifity is inversely correlated to the false positive rate. A low false positive rate is necessary because we expect that most differences found in a group of genomes will not be directly related to the convergent phenotypic change, but may come from neutral processes or be selected for reasons unrelated to the convergent phenotype (Bazykin *et al*., 2007; Rokas and Carroll, 2008; Zou and Zhang, 2015a). Therefore, among a large number of changes, only a small number will be associated with convergent phenotypic evolution. There will be very few positives to find, and a large number of negatives, which provides many opportunities for methods to predict false positives. To illustrate this point, we can use the numbers of substitutions inferred on terminal branches of the species tree provided in (Thomas and Hahn, 2015), based on transcriptome-wide analyses. If we take the example of microbats and dolphins, species that both evolved the ability to echolocate, (Thomas and Hahn, 2015) report roughly 4000 substitutions to different amino acids, which they call divergent, and 2000 substitutions to the exact same amino acid, which they call convergent, *i.e*. 6000 substitutions total. These numbers are in proportion with those reported in pairs of non-echolocating species, which was taken as evidence that the majority of the 2000 convergent substitutions detected by Parker *et al*. (2013) are not linked to the convergent evolution of echolocation. Instead they find that less than 7% of genes with convergent substitutions are also associated with positive selection, a number they choose as the true number of convergent substitutions. Based on these considerations, among the 6000 substitutions, 140 are truly convergent, and 5860 are not. If we were to apply a test that has a very respectable sensitivity of 98% and an equally good specificity of 98%, we would detect 0.98 ∗ 140 = 137 true positives, and 0.02 ∗ 5860 = 117 false positives. So, we would have a false discovery rate of 117*/*(117+137) = 46%, despite a test with excellent properties. We use these simple calculations later in the manuscript when presenting the results obtained with different methods.

The three methods to detect convergent evolution are as follow. The first method used in (Parker *et al*., 2013) is based on the comparison of two topologies, one for convergent sites, and the other for non-convergent sites. It is derived from earlier efforts by Castoe *et al*. (2009). Here, we named this method “Topological”. The second method used in (Foote *et al*., 2015; Thomas and Hahn, 2015; Zou and Zhang, 2015b) proposes to detect convergent changes related to a phenotypic change by focusing on substitutions to the exact same amino acid in each species with the convergent phenotype. We named this method “Identical”. Both methods can be used on rooted or unrooted trees, since they do not explicitly consider changes in the substitution models. Finally, the third method fleshes out our own definition of convergent shifts and is based on a modification of usual models of site evolution (Fig. 1). Under those models, any number of substitutions (including zero) can occur on a branch. To impose that convergent substitutions should occur on the branches where the phenotype changes, we introduce the `OneChange` model, shortened into OC, which imposes at least one substitution per site on the branch where it is applied. In addition to OC, we consider that convergent sites evolve according to different amino acid equilibrium frequencies (*i.e*. different profiles) in species with the ancestral or convergent phenotypes. Here, amino acid profiles are defined as profiles from (Si Quang *et al*., 2008) (see Fig. S1 in supplementary material), but other profiles could in principle be used. We named this model PCOC, for “Profile Change with One Change”, and also because it is the name of a beautiful bird.

**FIG. 1.**
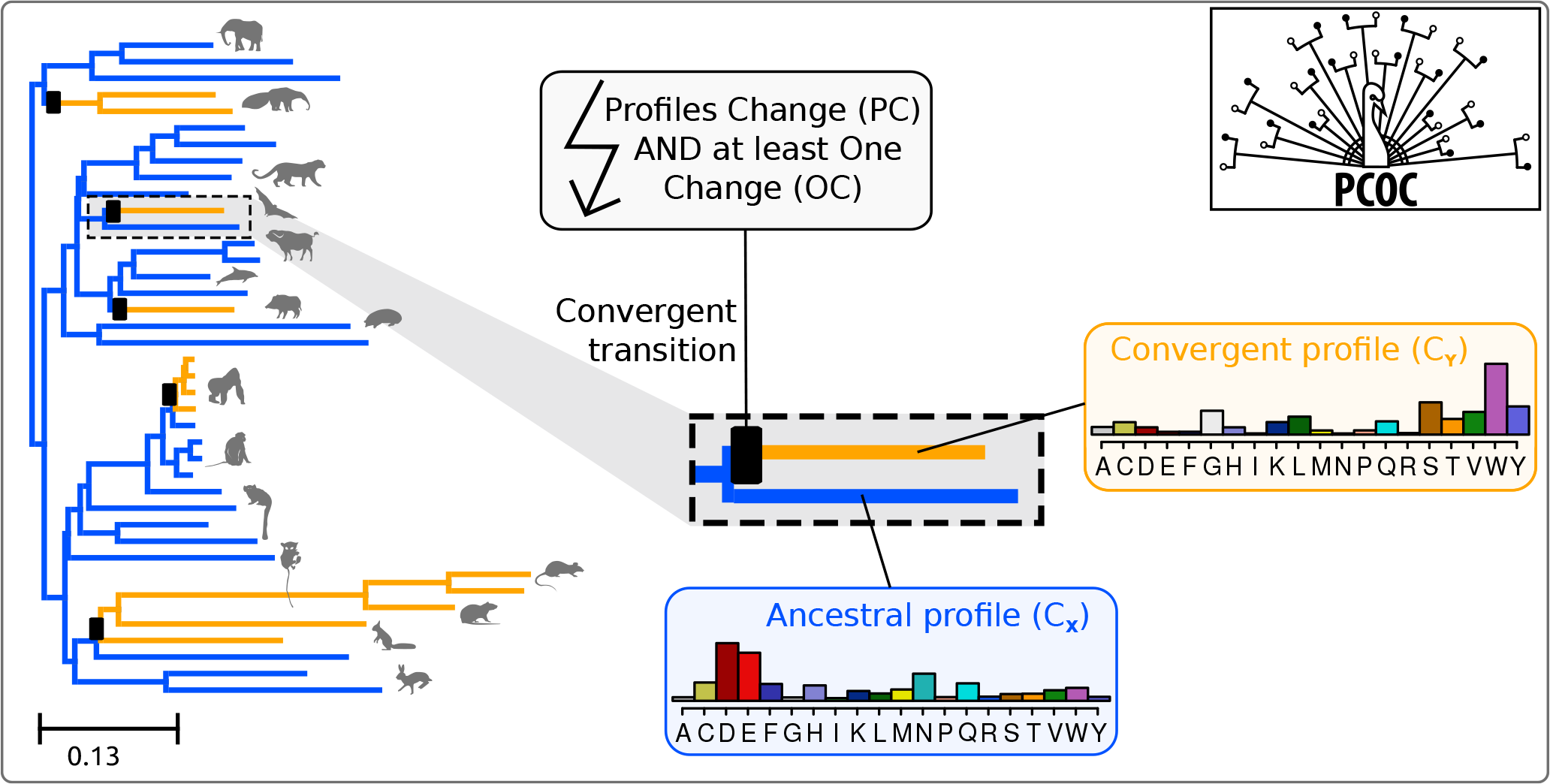
PCOC attempts to detect sites that are linked to the repeated evolution of a convergent phenotype. On the left, the Ensembl Mammalian phylogeny has been represented, and 5 transitions have been randomly placed on its branches (black boxes). On the branches with the boxes, PCOC imposes an amino acid profile change and the use of the OC model. The convergent profile is used in subsequent branches.

PCOC therefore combines two models, OC, which is new, and changes in amino acid profiles (PC), an idea that has been used before on single genes. In particular it has been used to study changes in selective constraints in the Influenza virus (Tamuri *et al*., 2009), or convergent evolution of a particular enzyme in C3/C4 plants (Studer *et al*., 2014). Recently such profile changing models have been extended into a Bayesian framework by Parto and Lartillot (Parto and Lartillot, 2017, 2018) for a gene-wise analysis of convergent evolution. In PCOC, it is possible to use only OC, or only PC, and in the manuscript we explore the properties of these two submodels PC and OC. PCOC detects convergent sites by comparing the fit of two models.

Under the convergent model, a site evolves under a commonly used model of protein evolution on most branches. Then, in clades with the convergent phenotype, it evolves under a model with a different vector of amino acid equilibrium frequencies. Further, we apply OC on branches where the phenotype has changed from ancestral to convergent, imposing that the model shift occurs at the beginning of the branch (but the substitution event can occur everywhere on this branch). As the PCOC model is by definition non-stationary, it requires a rooted tree. Under the non-convergent (null) model, a site evolves under a single amino acid profile throughout the phylogeny. We can thus compare the fit of the two models, the convergent and the non-convergent ones, on a given site of an alignment in terms of their likelihood to classify this site as convergent or non convergent. We implemented these models to perform sequence simulation as well as probabilistic inference in the Maximum Likelihood framework. Mathematical details are provided in the Methods section as well as in the supplementary material.

In this manuscript, we implement the PCOC model for simulation and estimation. We compare its efficiency to that of two existing methods for detecting convergent evolution and investigate its behaviour in a variety of conditions, changing the parameters of the simulation model, varying the number of convergent events, or introducing discrepancies between the simulation and inference conditions. Then we apply PCOC to a previously analyzed alignment of plant proteins where many convergent sites have been proposed. We find that although PCOC uses a different definition, it recovers many of the previously proposed convergent sites and conclude that this new model can be used on real data.

## Results

### Comparison of the three methods to detect convergent changes

We compared the performance of the Topological, Identical and PCOC approaches on simulations. We used empirical phylogenies, where a number of convergent transitions were placed randomly (from 2 to 7 events). In other simulations, we kept the empirical topologies, but fixed 5 convergent events and made branch length vary from small to large (Fig. 2). We have chosen thresholds that maximize the performance of the 3 methods to compare them fairly (see methods). However, the simulations are performed under our definition of convergent substitutions, which could advantage our method, fit for this definition, compared to the Topological and Identical methods. It is unclear how we could have avoided this bias. The Topological approach, with its operational definition, should be able to capture shifts in amino acid profiles, and could obtain very good results. The Identical approach is expected to have a much worse sensitivity, and can only capture convergent changes only when the convergent profile is very centered on a single amino acid. We will see that the results recover these broad tendencies. We used the mammalian subtree of the Ensembl Compara phylogeny, but similar results were obtained on other phylogenies (a phylogeny of birds from (Jarvis *et al*., 2014), a phylogeny of Rodents from (Schenk *et al*., 2013), and a phylogeny of the PEPC gene in sedges (Supplementary Fig. S17, S25 and S33)). PCOC outperforms the other approaches in the vast majority of conditions, by recovering higher proportions of true positives and lower proportions of false positives. Expectedly, PCOC and the Topological approaches both improve as the number of convergent changes increases (Fig. 2 A and B). However, the performance of the Identical method degrades as the number of changes increases, because it is rare that the exact same amino acid is found in e.g. 7 clades. As expected, the efficiency of all the methods increases as the distance between the simulated ancestral and convergent profiles increases (Supplementary Fig. S4). We also investigated the impact of the convergent profile itself, using a measure of its entropy. A profile with high entropy has similar frequencies for all 20 amino acids, whereas a profile with low entropy only has a few amino acids containing most of the probability mass. We find that PCOC is nearly insensitive to the entropy of the convergent profile, because its OC component itself is insensitive. However, both the identical and topological approaches have better results on convergent profiles with low entropy (Fig. S16). This result is expected for the Identical method, which should be best in cases where the probability mass of the convergent profile is all contained in one single amino acid.

**FIG. 2.**
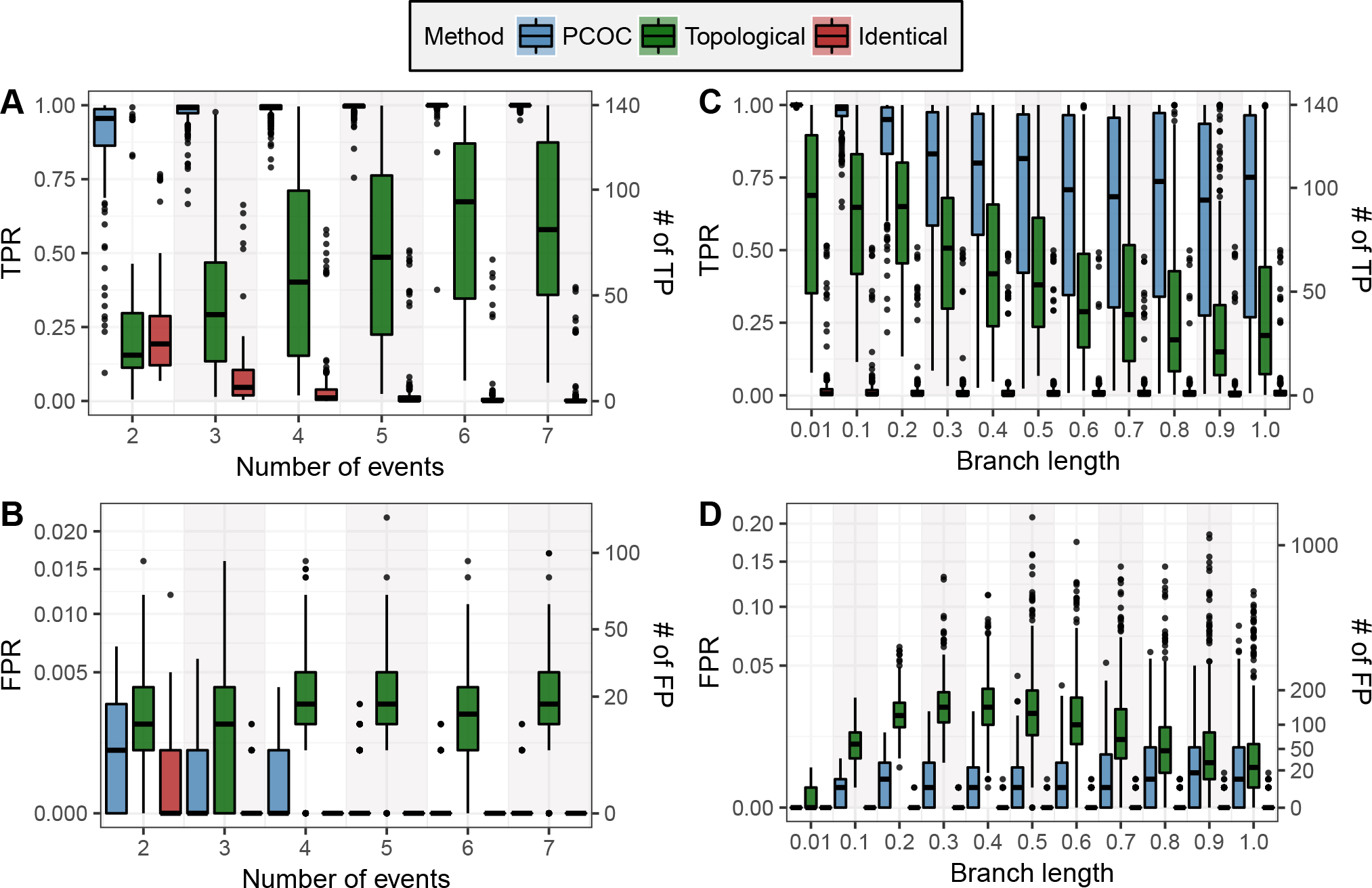
Comparison between the topological, identical and PCOC approaches to detect convergent substitutions. In A and B, we vary the number of convergent events from 2 to 7. In C and D, we set all branch lengths in the tree to a single value, ranging between 0.01 to 1.0 expected substitutions per site. The True Positive Rate (TPR) is the rate of TP among positives, *i.e*. the *sensitivity*, and the False Positive Rate (FPR) is the rate of FP among the negatives, *i.e*. 1*− specificity*. The right axes provide the numbers of true and false positives in the context of the example of the Introduction.

The performance of all methods tends to decrease as branch lengths become longer (Fig. 2, C and D). The Topological approach however predicts fewer false positives for branches nearing 1.0 expected substitution per site than for branches of length 0.5, but always performs worse than PCOC.

To ensure that PCOC was not unfairly favored in those tests, the above simulations have been performed using the C60 set of amino acid profile, while inference was performed using a different set of profiles, C10. We also tried to further complexify the simulations to make them harder for PCOC to analyze and evaluate how PCOC fares when some of its assumptions are violated. In particular, we used more than one amino-acid profile on the branches with the ancestral phenotype. To achieve this, we picked at random a few branches with the ancestral phenotype, and applied a different amino acid profile to those branches and the subsequent branches (Supplementary Fig. S8). We observed that PCOC’s performance did not change (Supplementary Fig. S9, S10). We also tested the performance of PCOC with mis-estimated branch lengths. To this end, we performed inferences on the trees used for simulation but after altering their branch lengths (see methods). The results did not seem to be affected by the amount of error introduced (Supplementary Fig. S11, S12).

We also assessed how PCOC was affected by misplacements of the events of convergent evolution. Fig. S13 shows that PCOC is more sensitive to the inclusion of a spurious event of convergent evolution than to the removal of an event of convergent evolution. However, PCOC still obtains better results than the topological or the identical approaches.

We also investigated how PCOC was affected by errors in the root of the tree by moving the root to neighboring branches of the root. Incorrect rooting did not seem to have much of an impact on PCOC (Fig. S15).

Finally, analyzing our set of random positioning of convergent transitions, we did not observe an influence of the proportion of leaves in convergent clades on the performance of the three methods (Supplementary Fig. S7). This differs from results obtained with the Identical method in (Thomas *et al*., 2017) which showed that fewer convergent sites were detected when more taxa with the convergent phenotype were used. However their experimental setup differs from ours in that we operate under a fixed total number of taxa whereas they changed the total number of taxa.

### PCOC’s performance draws on the PC and OC submodels

Fig. 3 shows the contributions of the PC and OC submodels to the performance of PCOC on the simulations with a single amino acid profile on ancestral branches. PCOC shows a much better performance than both its submodels. In most conditions, on those simulations, OC seems to perform better than PC. However we find that PC and OC perform best in different conditions. OC is most useful when branch lengths are short: in such conditions, encountering a substitution on a site provides a strong support for the OC model (Fig. 3 C and D). As soon as the expected number of substitutions approaches 0.5, the performance of OC drops markedly, because when a branch is longer than 0.5, a substitution is more likely than none, and then forcing one change on this branch has a minor impact on the transition probabilities. On the contrary, PC becomes more powerful as branch lengths increase, because PC can then exploit a larger number of substitutions both on branches with the ancestral profile and on branches with the convergent profile to identify a site as convergent. Similar results were obtained on three other phylogenies (Supplementary Fig. S18 to S39).

**FIG. 3.**
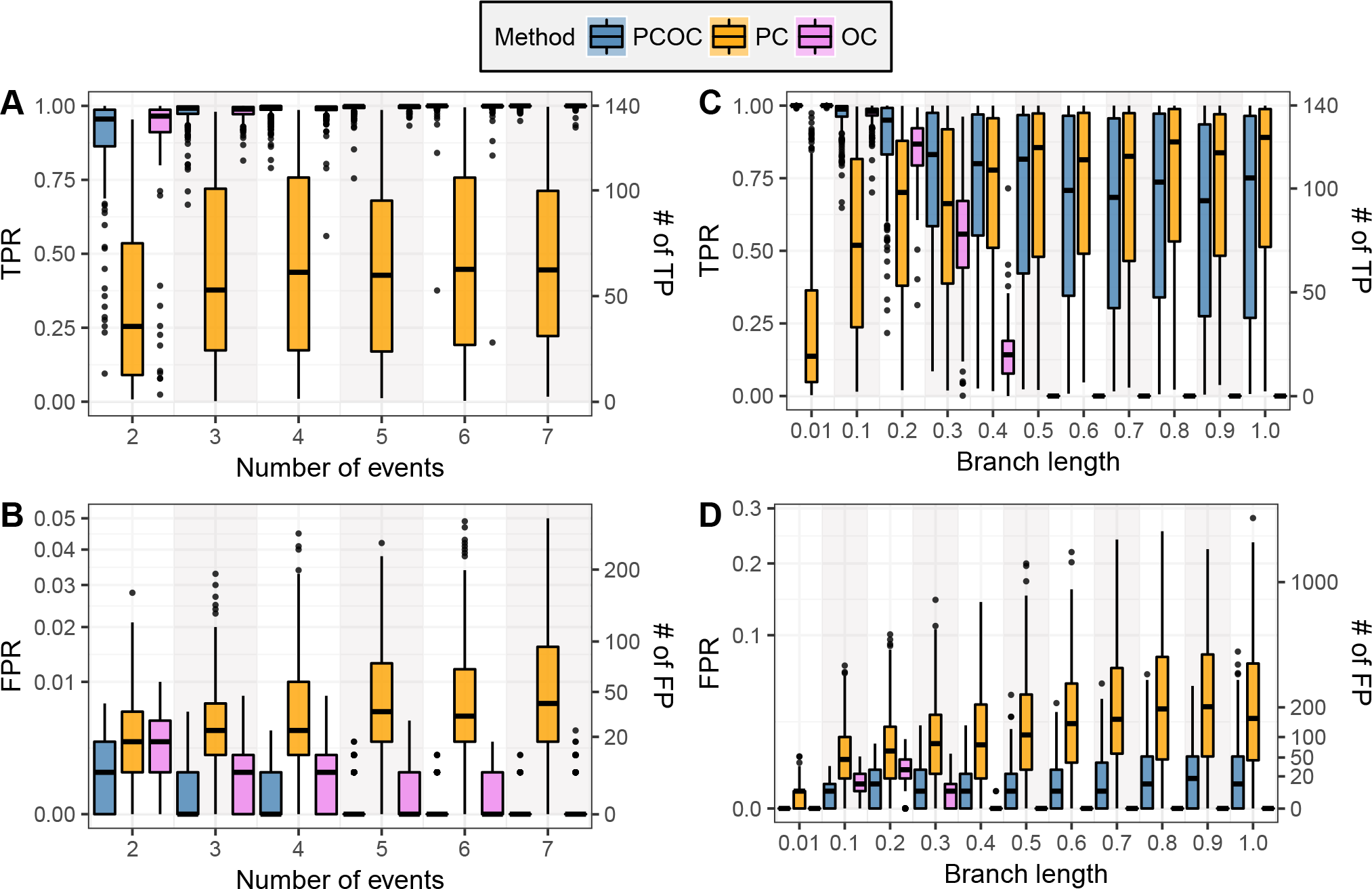
The power of PCOC draws upon its submodels PC and OC. See Fig. 2 for legend.

### Detection of convergent substitutions during repeated evolution of C4 metabolism in plants

Fig. 4 represents sites with predicted convergent substitutions in the PEPC protein occurring jointly with the transition towards C4 metabolism in sedges (Besnard *et al*., 2009). Sites are represented if they have been found convergent in (Besnard *et al*., 2009) (highlighted by a star), and/or by PCOC, using a threshold of 0.8. To detect convergent sites, Besnard *et al*. (2009) performed analyses of positive selection on the alignment, as well as comparative analyses with PEPC sequences from other plants. They proposed a set of 16 sites under positive selection (stars in Fig. 4). In addition to our analysis of the empirical alignment, we inferred convergent substitutions on simulations performed on the same topology, placing convergent transitions on the same branches, and using the C60 set of profiles to evaluate the numbers of false positives and negatives we should expect when running PCOC. In these simulations, with the same proportion of convergent sites as defined in the Introduction, we found that PCOC should produce neither false positives nor false negatives for an alignment of the same size as the empirical alignment. Accordingly, there is an important overlap between PCOC and the set of convergent sites proposed in (Besnard *et al*., 2009).

**FIG. 4.**
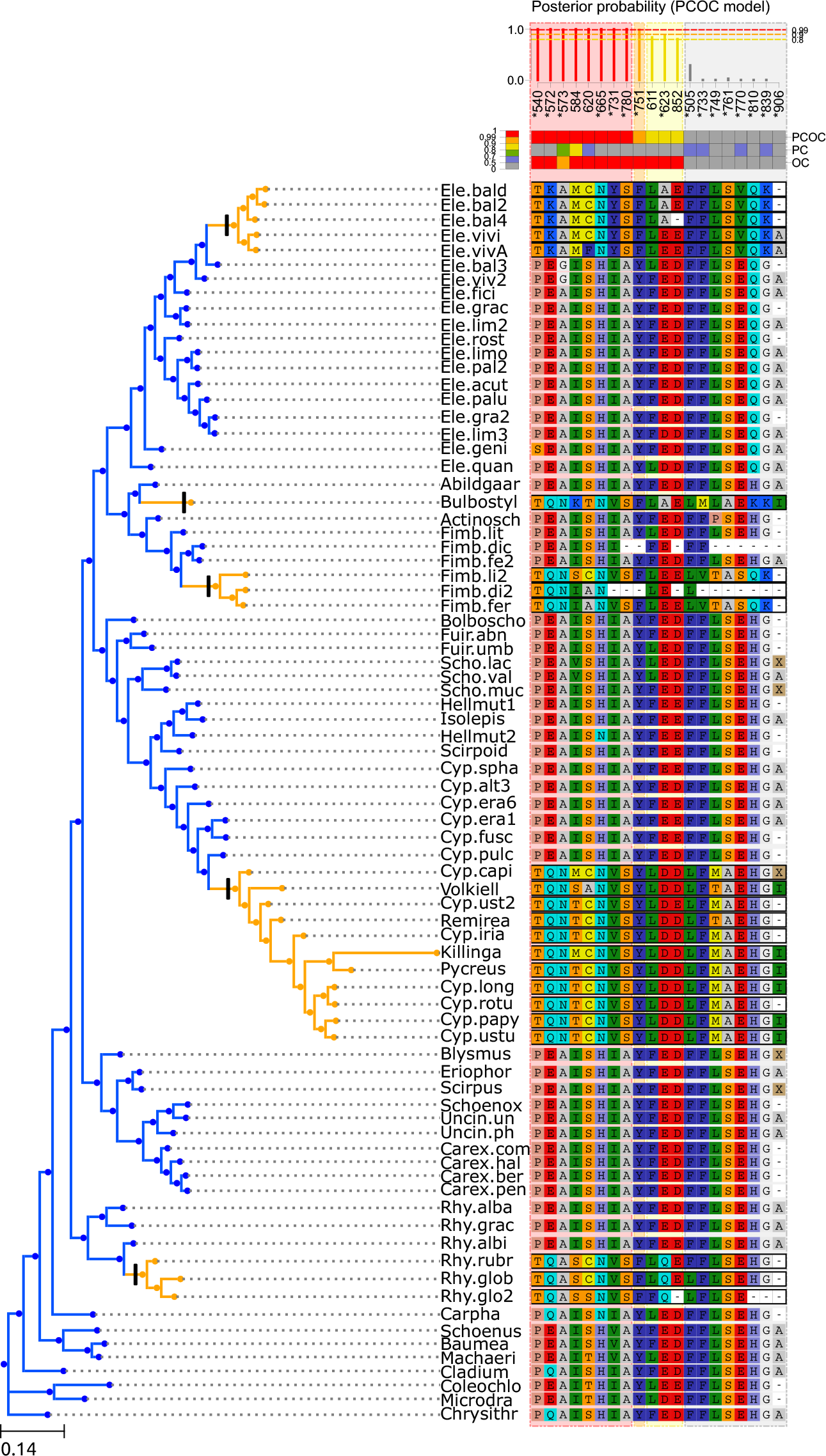
Detection of convergent substitutions using the PCOC toolkit in the PEPC protein in sedges. Sites are ordered by their posterior probability of being convergent according to the PCOC model. Only sites with a posterior probability (pp) according to the PCOC model above a given threshold (here, 0.8) or sites detected in (Besnard *et al*., 2009) (highlighted by a star) are represented. Sites are numbered according to Zea mays sequence (CAA33317) as in (Besnard *et al*., 2009). Posterior probabilities for the PCOC, PC, and OC models are summarized by colors, red for pp ≥ 0.99, orange for pp ≥ 0.9, yellow for pp *≥* 0.8 and gray for pp *<* 0.8.

Their intersection contains 8 sites (both with a star and in red, orange or yellow on the top of Fig. 4), and their union 20 sites. Only four sites predicted by PCOC have not been proposed in (Besnard *et al*., 2009). Further, manual inspection of the two new sites with the best posterior probabilities (positions 584, 620) suggests that they have undergone substitutions inside each of the C4 clades, possibly on the branch ancestral to those clades, and towards amino acids that are seldom found in the gene sequences from C3 species. To better understand why PCOC detects these two sites, we looked at the separate posterior probability of the PC and OC models for each of those two sites. In both cases, the very high posterior probability of PCOC is due in large part to the support for OC (pp*>*0.99), but the support for PC is also superior to 0.5 (0.82 and 0.66 for positions 584 and 620 respectively). The two other sites with lower posterior probabilities (611 and 852) are not as convincing, and are identified only thanks to the OC component of PCOC. In addition, there are 8 positions classified only by Besnard *et al*. (2009) as convergent and not predicted as convergent by PCOC, because they each underwent substitutions only in a subset of the C4 clades out of 5: 4 for position 505, 3 for position 761, 839, 2 for positions 749, 770, 810 and 906 and 1 for position 733. For all those sites, there is no support for OC and at best weak support for PC, because those sites do not fit PCOC’s definition of a convergent site.

We also performed analyses by using only the OC and PC submodels. PC only predicts 7 sites as convergent (Supplementary Fig. S41), and none of them are predicted in (Besnard *et al*., 2009). Among the 14 sites it predicts as convergent (Supplementary Fig. S42), OC finds 8 sites also predicted by Besnard *et al*. (2009), like PCOC. The similarity between the sites selected by OC and those selected by PCOC is large, but two sites, sites 518 and 579, are predicted as convergent by OC but not by PCOC, and are not found in (Besnard *et al*., 2009). Overall, PCOC’s predictions appear to be derived mostly from the OC submodel rather than from the PC submodel, and are consistent with a previously published detailed analysis of an amino acid alignment. New positions suggested by PCOC represent candidates for convergent substitutions.

## Discussion

### Defining convergent amino-acid evolution

In this work we have used a new definition of convergent events of genomic evolution, focusing on events that involve single amino acid substitutions that occur simultaneously (at the scale of single branches) with convergent phenotypic changes. This definition fits causative changes, or changes so intimately associated to the convergent phenotype that they occur very shortly after the phenotype has changed. We developed PCOC to simulate and detect changes according to this definition.

### PCOC accurately detects events of convergent amino-acid evolution

Compared to two previously proposed methods to detect convergent substitutions, PCOC has best power to detect changes that fit its definition. However, because PCOC relies on two submodels PC and OC, in principle it can also capture convergent changes that do not perfectly fit the definition above (Fig. 3). For instance, it may be able to detect substitutions that occur systematically on branches where the phenotype changed, irrespective of whether this was associated to a profile change, thanks to the OC component of PCOC. OC may thus recover sites detected by methods that look for accelerations on branches where the phenotypes changed (Partha *et al*., 2017). Similarly, thanks to its PC component, it may be able to detect sites that have not undergone substitutions on the branches where the phenotype changed, but that have undergone substitutions in underlying branches according to the convergent amino acid profile.

In practice, the PC submodel does not seem to contribute as much as the OC submodel, as seen from the C4 convergence example (Figs. 4, Sup. Fig. S41 and S42). It is unclear whether this is an inherent limitation of the data set, where branch lengths are at most 0.217, of the PC approach, or if better fitting profiles could improve PC’s performance. Regarding branch lengths, PC could indeed contribute more than OC to PCOC on data sets where branch lengths are long (Fig. S6). Regarding better fitting profiles, inferences performed under the same C60 model as that used for simulation show that the PC component is still minor compared to the OC component (Fig. S5), even when the profiles perfectly fit the simulation. However, more pointy profiles, where only a few amino acids have non-zero probability, may fit the data better. Such profiles are uncommon in the C60 and C10 sets, but they would better correspond to the particular subset of amino acids found at a given site in the convergent species.

### Comparison between PCOC and mutation-selection models

Parto and Lartillot (Parto and Lartillot, 2017, 2018) have used a mutation-selection model to detect convergent evolution in single gene sequences. Mutation-selection models are codon models that attempt to distinguish the contribution of the mutational process at the DNA level from the contribution of the selection process, typically at the amino acid level. PCOC is a model of amino acid sequence evolution and therefore ignores phenomena that happen at the DNA level. In both PCOC and mutation-selection models, convergence is expected to be linked to changes in amino acid profiles; in fact, the PC submodel of PCOC can be thought of as an approximation of Parto and Lartillot’s model, in the Maximum Likelihood framework, with a fixed set of profiles. However PCOC further adds the OC submodel, which enables it to detect repeated accelerations of the evolution of a site on the branches where the phenotype changed, even in the absence of a profile change. Further, PCOC benefits from a speed advantage over mutation-selection models as implemented in (Parto and Lartillot, 2017, 2018) for two reasons. First, because it works with protein sequences instead of codon sequences, which reduces the time required to compute the likelihood of a model. Second, because PCOC does not attempt to estimate amino acid profiles: instead it draws from profiles that have been estimated from large numbers of alignments. For these reasons PCOC can be used easily at the scale of whole genomes (see below).

### PCOC is a tool to simulate and detect convergent genomic evolution

We developed PCOC as a set of tools to perform simulation and detection of convergent evolution in sequences. These tools are user-friendly and require a gene tree provided by the user. It takes about 40 seconds to run the detection tool on a laptop for a data set with 79 leaves and 458 sites with the C10 set of profiles, and up to 20 minutes with the C60 set of profiles. The PCOC tool-kit is open source and available on GitHub https://github.com/CarineRey/pcoc with a tutorial. Simulations can be used to test the capacity of PCOC or other methods to detect convergent evolution on a specific data set, with its idiosyncratic characteristics. We have observed that the power of the methods depends on the number of independent convergent phenotypic changes, on branch lengths, and on the tree topology. These simulations can also be used to choose thresholds for controlling the amounts of false positives and false negatives. It is also easy to simulate sites with and without convergent evolution, for testing other methods.

### Using PCOC with genomic data

We have not attempted to work at the level of entire gene sequences or even functional groups of genes, whereby the evidence obtained at the level of individual sites would be used collectively over the entire gene length or over several genes with a particular function to classify a gene or group of genes as convergent or not. However, other works have developed methods to work above the level of single sites (Chabrol *et al*., 2017; Marcovitz *et al*., 2017), and our method is compatible with these. Both these approaches detect convergent substitutions that fit the definition of (Foote *et al*., 2015; Zhang and Kumar, 1997), but use different approaches to classify genes as convergent or not. Chabrol *et al*. (2017) combine their site-wise analysis with a procedure involving simulations according to a null model to classify genes as convergent or not. This simulation procedure is easy to perform with the PCOC toolkit. In particular, to investigate convergent evolution in a gene, we suggest that first convergent sites are identified using PCOC. Then, using the same tree and same parameters that were used for detection, one would perform simulations of a large number of sites with convergent evolution, and of sites without convergent evolution. PCOC would then be run on those simulated sites, which would provide the amount of true positives and false negatives. Such an approach can be used to assess the false discovery rate associated with the selection of candidate convergent sites in the empirical data. We applied this approach in our study of the C3/C4 alignment and described the procedure in the PCOC tutorial.

### Possible improvements to PCOC

PCOC relies on a set of profiles empirically built from a large number of alignments (Si Quang *et al*., 2008). These profiles were constructed to accurately model protein evolution in a time-homogeneous manner, and may be suboptimal for describing the evolution of sites that switch between two distinct profiles. Other profiles could be used although this has not yet been implemented in PCOC.

PCOC relies on a more general definition of convergent genomic events than the usual definition involving substitutions to a specific amino acid, but still does not account for other types of convergent events. For instance, PCOC has not been designed to deal with convergent relaxations of selection. To do this, in (Marcovitz *et al*., 2017), groups of candidate genes that contain an excess of convergent substitutions are filtered using divergent substitutions, *i.e*. substitutions to different amino acids in the convergent species. PCOC does not rely on the definition of (Foote *et al*., 2015; Zhang and Kumar, 1997), and therefore it is uneasy to define such divergent substitutions. In our case, to distinguish convergent relaxations, we would rely on the fact that substitutions should accumulate in the convergent branches, but no particular profile of amino-acids should be favored. For example, this corresponds to a shift from a pointy to a broad amino-acid profile. Detecting this requires to access the scores for all profiles in PCOC, and contrast their pointedness. This is not yet implemented in PCOC. To detect potential cases of convergent relaxations, we could also filter candidate genes based on branch lengths in convergent species: genes under relaxed selection specifically in lineages with the convergent phenotype are expected to have longer branches in those lineages.

Finally, the requirement linked to the OC submodel that convergent sites should undergo substitutions simultaneously with each convergent transition may be too strict: in some cases it will be sufficient to consider a site as convergent if it undergoes substitutions on a large subset of those transitions. PCOC could be modified to fit such situations by using a mixture model, so that according to a probability *p* the OC submodel would be used on the branches subtending convergent clades, and according to 1*−p* the OC submodel would not be used. The estimation of this single parameter *p* would probably not incur an important computational cost.

## Materials and Methods

### A new probabilistic model of convergent evolution

We adopt a biochemical point of view and consider that adaptive convergence drives the preference at a given site towards amino acids that share specific properties. We do not define those properties *a priori*, but instead consider a set of amino acid profiles, empirically built from a large number of alignments (Si Quang *et al*., 2008). These profiles serve as a proxy to amino acid fitnesses at a given site: more frequent amino acids in the profiles have higher stationary frequencies, as in mutation-selection models (Parto and Lartillot, 2017). Following this Profile Change (PC) model, a convergent site will exhibit a preference in all convergent clades towards a specific profile, different from an ancestral profile, whereas a non-convergent site will remain with the same profile in all the tree. In our simulations, we also consider the possibility that a non-convergent site alternates randomly between a few different profiles along the phylogeny on branches with the ancestral phenotype, but switches to a particular single profile on branches with the convergent phenotype. In addition, we consider that a substitution must occur when a convergent site switches from the ancestral profile to the convergent profile, and to this end we implemented the OneChange (OC) model. The combination of PC and OC into PCOC models the situation where the convergent phenotype is tightly linked to a given type of amino acid at a certain position, so much so that it can be considered necessary or at least highly advantageous for the phenotype to have one of the fittest amino acids from the convergent profile at this position. Our approach therefore does not attempt to model positions that change to a convergent amino acid profile after the switch from the ancestral to the convergent phenotype has occurred, and which would be non-causative substitutions. Such sites would be appropriately modeled by PC alone, but not quite as well by PCOC.

### PCOC Tool-kit: a tool for simulation and inference of convergent substitutions

#### Simulation process

To evaluate the ability of detection methods to detect convergent sites, we performed two types of simulation. In one type, we simulate under convergent evolution, varying the parameters of the evolutionary model (e.g. varying the number of convergent transitions). This allows us to estimate the sensitivity of the methods. In the other type we simulate without any event of convergent evolution. This allows us to assess the specificity of the methods. In each case, we simulated 1000 sites. To simulate convergent evolution, we aimed at placing events of convergent evolution uniformly on a species tree, irrespective of branch length. We were interested in the impact of the number of events of convergent evolution on our power to detect it and placed between 2 and 7 events. To avoid any bias in the location of these events, in all cases we drew uniformly exactly 7 potential events, so that all events were in independent clades. From these 7 events we then subsampled the desired number of events of convergence. All branches in the clades below those events were labeled “convergent”, and all other branches (above these events and in the non-convergent clades) labeled “ancestral”. A particular amino acid fitness profile *c_x_* was used for ancestral branches, another *c_y_* for convergent branches and we applied the OneChange model with the *c_y_* profile on the branch where the switch to the convergent phenotype was positioned. The switch was placed at the very beginning of the branch. We randomly drew amino acid profiles from the C60 model (Si Quang *et al*., 2008) (Supplementary Fig. S1) and did not attempt to test all pairs of C60 profiles in order to save computation time and slightly reduce our carbon footprint. We also performed additional simulations where more than one profile was used on branches with the ancestral phenotype (Supplementary Fig. S8, S9 and S10). Although C60 was built to describe amino acid sequence evolution in a time-homogeneous manner, we assume that this limited set of profiles provides a rough approximation to the set of possible amino acid profiles. In addition to the simulations with convergent events that we used to measure the proportion of True Positives (TP) and False Negatives (FN) of the methods, we performed similar simulations (*i.e*. using the same trees) where the ancestral profile is used for all branches of the phylogeny, to measure their proportion of True Negative (TN) and False Positive (FP).

Sequence evolution was simulated along the phylogenetic tree using the model associated to each branch, with rate heterogeneity across sites according to a Gamma distribution discretized in 4 classes (Yang, 1994) with the *α* parameter set to 1.0, using bppseqgen (Dutheil and Boussau, 2008).

#### Inference methods

For each of the three compared approaches, we have to infer if a site is convergent.

For the PCOC, PC, OC and the Topological methods, the decision is controlled by a threshold on the *a posteriori* probability of the convergent model vs the null model, using a uniform prior. We used bppml (Dutheil and Boussau, 2008) to measure the likelihood of each model.

To compare the studied methods fairly, we tuned this threshold for each method to reach its optimal performance. We use the Matthews correlation coefficient (MCC) (Matthews, 1975) as a measure of the performance because the MCC takes into account the proportions of positives and negatives which are expected to be heavily biased in our case as we saw in the Introduction. Therefore we chose the threshold so as to maximize the MCC of each method using the proportions of the Introduction example. (Supplementary Fig. S2).

Below we describe the procedure we adopted to call a site as convergent for each of the three compared approaches.

##### • PCOC approach

In accordance with our definition of convergence and our simulation procedure, we used a model-based inference to detect convergent substitutions. We used the branch lengths that had been used for simulation for inference, but we checked that the impact of errors in branch lengths on inference was minimal (Fig. S11 and S12). We used the C10 set of profiles from the CAT model (Si Quang *et al*., 2008), containing 10 profiles, to be in a more realistic scenario where the CAT profiles used in the simulation (C60) are not those used for inference. However we checked that using the same C60 set of profiles for inference and simulation yielded very similar results (Fig. S5). For each i in {1..10} and for each j in {1..10} such as *i ≠ j*, we calculated the likelihood of two models: one, *M*0*_i_*, in which the same profile *c_i_* is used on all branches, and another model,*M*1*_i/j_*, in which the profile *ci* is used only on “ancestral” branches, and the profile *c_j_* on “convergent” branches. We explain in details how one can compute the likelihood under *M*1 in the supplementary material, section 2. Then, we compared the likelihoods of two average models, *M*0 and *M*1. The likelihood of *M*0 is computed as the mean of the likelihoods of the *M*0*_i_* models and the likelihood of *M* 1 as the mean of the likelihoods of the *M* 1*_i/j_* models.

We classified each site as a positive or a negative using an Empirical Bayes approach. A positive is a site predicted to have evolved according to the heterogeneous model *M*1, and a negative according to the homogeneous model *M*0. For each site *i*, we computed the likelihood of the *M*1 model *P* (*si|M*1) and of *M*0 *P* (*si|M* 0). We computed the empirical posterior probability of *M*1 with a uniform prior on each model: *P*(*M*1*|si*)= *P*(*si|M*1)*/*(*P*(*si|M*1)+*P* (*si|M* 0)). A positive is defined such that *P* (*M*1*|si*) *>* 0.99 for the PCOC and the OC models and 0.9 for the PC model.

##### • Topological approach

We also performed comparisons of likelihoods with two different topologies, as in (Parker *et al*., 2013). The rationale of this approach is that, for sites showing convergence, the phylogenetic signal would prefer to cluster together convergent branches. So, for these sites, the true tree should be less likely than the tree for which the convergent branches are together, named “convergent tree”. We present in Supplementary Material the algorithm we used to construct convergent trees and an example of such a “convergent tree” (Supplementary Fig. S3).

We computed for each site, the mean of the likelihoods with the ancestral model *ci* applied on all branches for each i in {1..10} for the true and the convergent trees. And, as in the method based on heterogeneous models, we considered a site as convergent when the empirical posterior probability of the convergent tree was above 0.9.

##### • Approach based on ancestral reconstruction

To detect convergent substitutions as in (Foote *et al*., 2015; Thomas and Hahn, 2015; Zou and Zhang, 2015b), we considered the branches ancestral to convergent clades.

We declared a substitution on a given site as convergent if all substitutions on the ancestral branches were towards the exact same amino acid.

#### Statistical measures of the performance

Finally, we measured the power of the three methods of detection on simulations using their specificity, sensitivity, and MCC (Supplementary Fig. S4, S6, S7, S9, S10, S11, S12, S18 to S24, S26 to S32 and S34 to S40).

### Simulations to assess the impact of the number of convergent transitions

We used the simulator and benchmark tool of the PCOC toolkit to produce the data used in the panels A and B of Fig. 2 and 3. We extracted the subtree containing mammals only from the Ensembl Compara tree (Herrero *et al*., 2016; Yates *et al*., 2016), and used it to position a random number X of convergent events between 2 and 7. We repeated this procedure 160 times. For each random assignment of convergent events, we sampled 10 pairs of C60 profiles and for each pair simulated 1000 convergent sites using both profiles and 1000 non-convergent sites using only the ancestral profile.

### Simulations to assess the impact of branch lengths

We used the simulator and benchmark tool of the PCOC toolkit to produce the data used in the panels C and D of Fig. 2 and 3. We used the same tree as above, and set all its branch lengths to values between 0.01 and 1. For each branch length value, we performed 32 replicates by randomly placing 5 events of convergent evolution in the phylogeny. For each random assignment of convergent events, we simulated alignments with 10 pairs of C60 profiles and for each pair simulated 1000 convergent sites using both profiles and 1000 non-convergent sites using only the ancestral profile.

### PCOC Tool-kit: Detector tool, test on real data

We used the detector tool of the PCOC toolkit to build Fig. 4. It takes about 40 seconds to run on a laptop for a data set with 79 leaves and 458 sites with the C10 set of profiles, and up to 20 minutes with the C60 set of profiles. The nucleotide alignment and tree topology come from (Besnard *et al*., 2009). As the detector tool of the PCOC toolkit needs a tree and an amino-acid alignment, we inferred branch lengths on the fixed topology using phyml (Guindon *et al*., 2010) with the GTR model using the nucleotide alignment and obtained the amino-acid alignment by translating the nucleotide sequences. For clarity, we only showed sites if they had a posterior probability above 0.8 according to the PCOC model (See Supplementary Fig. S41 and S42 for the PC and OC models).

## Conclusion

We have proposed a new definition of convergent substitutions that contains and relaxes the commonly used definition from (Zhang and Kumar, 1997). We have implemented a model embodying this definition into simulation and inference methods, and find that our method has better power to detect convergent changes than previously proposed approaches. It is sufficiently fast to be applied on large data sets, and should be useful to detect traces of convergent sequence evolution on genome-scale data sets.

## Supplementary Materials

Supplementary materials are available at Molecular Biology and Evolution online (http://www.mbe.oxfordjournals.org/).

## Acknowledgments

This work was performed using the computing facilities of the CC LBBEPRABI. We thank Thibault Lorin, Vincent Lanore, Gilles Didier, Philippe Veber, Nicolas Lartillot and Vincent Daubin for fruitful discussions. Fundings: ANR-15-CE32-0005 “Convergenomix”, ANR-10-BINF-01-01 “Ancestrome”, ANR-11-JSV6-00501 “Convergdent”. We thank Pauline S´emon for the PCOC logo. The work presented in this manuscript involved more than 400 computer.days.

